# Direct visualization of translational GTPase factor-pool formed around the archaeal ribosomal P-stalk by high-speed atomic force microscopy

**DOI:** 10.1101/2020.06.30.176701

**Authors:** Hirotatsu Imai, Toshio Uchiumi, Noriyuki Kodera

## Abstract

The ribosomal stalk protein plays an essential role in the recruitment of translational GTPase factors EF1A and EF2 to the ribosome and their GTP hydrolysis for efficient translation elongation. However, due to the flexible nature of the ribosomal stalk, its structural dynamics and mechanism of action remain unclear. Here, we applied high-speed atomic force microscopy (HS-AFM) to directly visualize the action of the archaeal ribosomal stalk (P-stalk). HS-AFM movies clearly demonstrated the wobbling motion of the P-stalk on the large ribosomal subunit, where the stalk base adopted two conformational states, a predicted canonical state, and a newly identified flipped state. Intriguingly, archaeal aEF1A and aEF2 molecules spontaneously assembled around the ribosomal P-stalk up to the maximum number of available binding sites. These results provide the first visual evidence for the factor-pooling mechanism and reveal that the ribosomal P-stalk promotes translation elongation by increasing the local concentration of translational GTPase factors.

## Introduction

The ribosome translates genetic information by interacting with various translational GTPase factors (trGTPases) in each step of translation (initiation, elongation, termination, and recycling). Translation elongation is mediated by two trGTPases: EF1A (EF-Tu in bacteria), which delivers aminoacyl-tRNA (aa-tRNA) to the ribosomal A-site; and EF2 (EF-G in bacteria), which catalyzes translocation (1, 2). EF1A and EF2 alternately bind to a common binding site, the factor-binding center of the ribosome, and dissociate upon GTP hydrolysis. In addition, EF1A has to continuously deliver aa-tRNA to the A-site until the ribosome accepts the cognate aa-tRNA. In order to efficiently repeat these EF1A-and EF2-dependent steps in accurate translation elongation, ribosomes possess a delicate molecular equipment for binding and dissociation of EF1A and EF2.

A set of ribosomal proteins, which is called the “stalk”, plays a crucial role in the recruitment of trGTPases to the factor-binding center and the activation of subsequent GTP hydrolysis (3–5). Previous biochemical studies revealed that the *E. coli* ribosomal L7/L12-stalk allows the association of the EF-Tu•GTP•aa-tRNA ternary complex and EF-G•GTP with the ribosome to take place more rapidly than would be expected by free diffusion (6, 7). The ribosomal stalk is conserved in all domains of life and present in the large subunit of the ribosomes. In archaea, one copy of the ribosomal protein aP0 binds directly to the 23S ribosomal RNA, and three homodimers of the stalk protein aP1 form the heptameric aP0•(aP1•aP1)_3_ P-stalk complex (7, 8). In eukaryotes, the ribosomal protein P0 associates with two P1•P2 heterodimers to form the P0•(P1•P2)_2_ P-stalk complex, while the bacterial ribosomal L7/L12-stalk are formed by the association of bL10 and two or three bL12 homodimers (4, 10). aP1 consists of three domains: the N-terminal dimerization domain which is also required for binding to aP0, the flexible hinge, and the C-terminal region which directly binds to each trGTPase (11–14). A C-terminal region homologous to aP1 is also present in aP0, and thus the archaeal aP0•(aP1•aP1)_3_ P-stalk has seven binding sites for the trGTPases. Sequence comparison shows that the archaeal aP1 and eukaryotic P1/P2 stalk proteins are homologous to each other. However, aP1 and P1/P2 show little similarity to the bacterial stalk protein bL12 (15), although they play a common role in mediating the efficient turnover of trGTPases.

Isolated ribosomal stalks have been well characterized by biochemical and structural analyses, whereas their mechanism of action and structure-function relationship in the ribosome-bound states are poorly understood. Although early electron micrographs identified a stalk-like protrusion composed of bL12 on the *E. coli* 50S subunit (16–18), in recent structural studies, such as X-ray crystallography and cryo-electron microscopy, the overall structure of the ribosomal stalk has not been detected due to its flexible nature (19–23). To date, only part of the C-terminal domain of bL12, which bound to EF-Tu or EF-G on the factor-binding center, has been visualized in several structural studies (24–29). Therefore, our knowledge about the binding dynamics between free trGTPases and their multiple binding sites on the ribosomal stalk is extremely limited.

At least two mechanistic functions of the ribosomal stalk have been discussed (5). One is the factor-pooling function, in which the multiple arms of the ribosomal stalk recruit trGTPases to the factor-binding center by binding and thereby increasing the local concentration of trGTPases (4, 9). The other is an ability to bind and stabilize trGTPases associated with the sarcin/ricin loop of 23/28S rRNA, which catalyzes GTP hydrolysis (30). Importantly, these two functions are not mutually exclusive, and both mechanisms presumably contribute to efficient recruitment and turnover of trGTPases on the ribosome. However, conclusive evidence has not been obtained for either case.

Here, we performed single-molecule observations using high-speed atomic force microscopy (HS-AFM) to investigate the structural dynamics of the ribosomal stalk and its binding to trGTPases. HS-AFM clearly visualized the flexible structure of the archaeal P-stalk base on the 50S subunit. Furthermore, we observed that multiple archaeal EF2 (aEF2) molecules were localized around the factor-binding center in the ribosomal P-stalk dependent manner, and this preferential distribution was similarly observed for archaeal EF1A (aEF1A). These results reveal that the ribosomal stalk recruits the elongation factors to pool around itself while flexibly moving on the ribosome. These mechanistic behaviors are consistent with a factor-pooling mechanism that contributes to efficient binding and action of the elongation factors.

## Results

### Observation of the ribosomal stalk on the large subunit by HS-AFM

We first determined the optimal conditions for immobilization of the ribosomes to a mica surface, an atomically flat substrate that makes an ideal sample stage for HS-AFM. The 50S and 30S ribosomal subunits from *Escherichia coli* were gently immobilized on (3-aminopropyl)triethoxysilane-treated mica (AP-mica) (Supplementary Fig. 1a–e). In this condition, the heights of the 50S and 30S were estimated to be ∼16 nm and ∼12 nm, respectively (Supplementary Fig. 1f, g). Given the estimated height of the ribosome, and the fact that AP-mica carries a positive surface charge, it is highly likely that the ribosomal subunits are immobilized to the stage via their inter-subunit rRNA surface which is negatively charged (Supplementary Fig. 1h, i). In HS-AFM images of the bacterial 50S ribosomal subunit, a small protrusion that seems to be the bL12 stalk base was detected (Supplementary Fig. 1a, b, d, Supplementary Movie 1). Unfortunately, no further clear images were obtained. Sequence alignments and structural studies show that the archaeal ribosomal P-stalk is larger than that of other kingdoms (Supplementary Fig. 2). Therefore, we targeted the 50S subunit from a hyper-thermophilic archaeon *Pyrococcus furiosus* to investigate the structural dynamics of the ribosomal stalk. We purified the 50S subunit from archaeal culture, and we succeeded in obtaining a clear HS-AFM image of the 50S subunit which displays three protrusions (Fig. 1a). The structural model of the archaeal 50S subunit and line profile analysis suggest that these three protuberances presumably represent the L1 stalk, the central protuberance (CP), and the P-stalk base including the C-terminal part of aP0 (SB), respectively (Fig. 1b).

**Figure 1.**
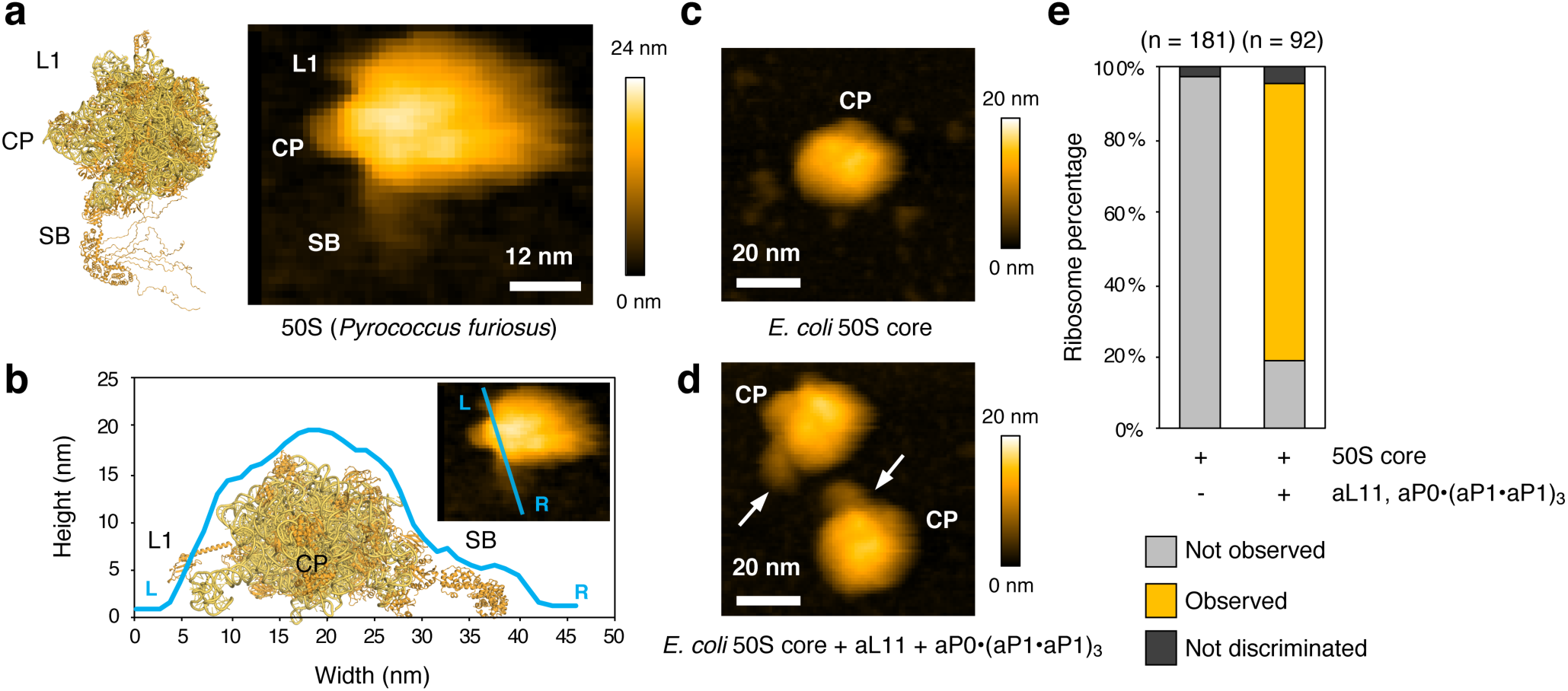
HS-AFM images of the archaeal ribosomal stalk on 50S ribosomal subunit. **a**, Left, a structural docking model of 50S subunit from *P. furiosus* and aP0•(aP1•aP1)_3_ P-stalk complex made from two coordinates [PDB code: 3A1Y and 4V6U]. The hinge regions and C-terminal regions of aP0 and aP1 are modeled arbitrarily. Right, a HS-AFM image of 50S subunit on AP-mica surface (L1: L1 stalk, CP: central protuberance, SB: P-stalk base). Scan area, 60 × 50 nm with 50 × 40 pixels; scan speed, 150 ms/frame; scale bar, 12 nm. **b**, Cross-sectional profile along the cyan-line drawn in the HS-AFM image in **a. c, d**, HS-AFM images of the 50S core (**c**) or the hybrid 50S (**d**) on AP-mica surface. Scan area, 80 × 80 nm with 80 × 80 pixels; scan speed, 250 ms/frame; scale bar, 20 nm. White arrows indicate the ribosomal P-stalk. **e**, Percentage of 50S subunits in which the ribosomal stalk was observed, related to **d** (Gray: not observed, orange: observed, black: not discriminated).

**Figure 2.**
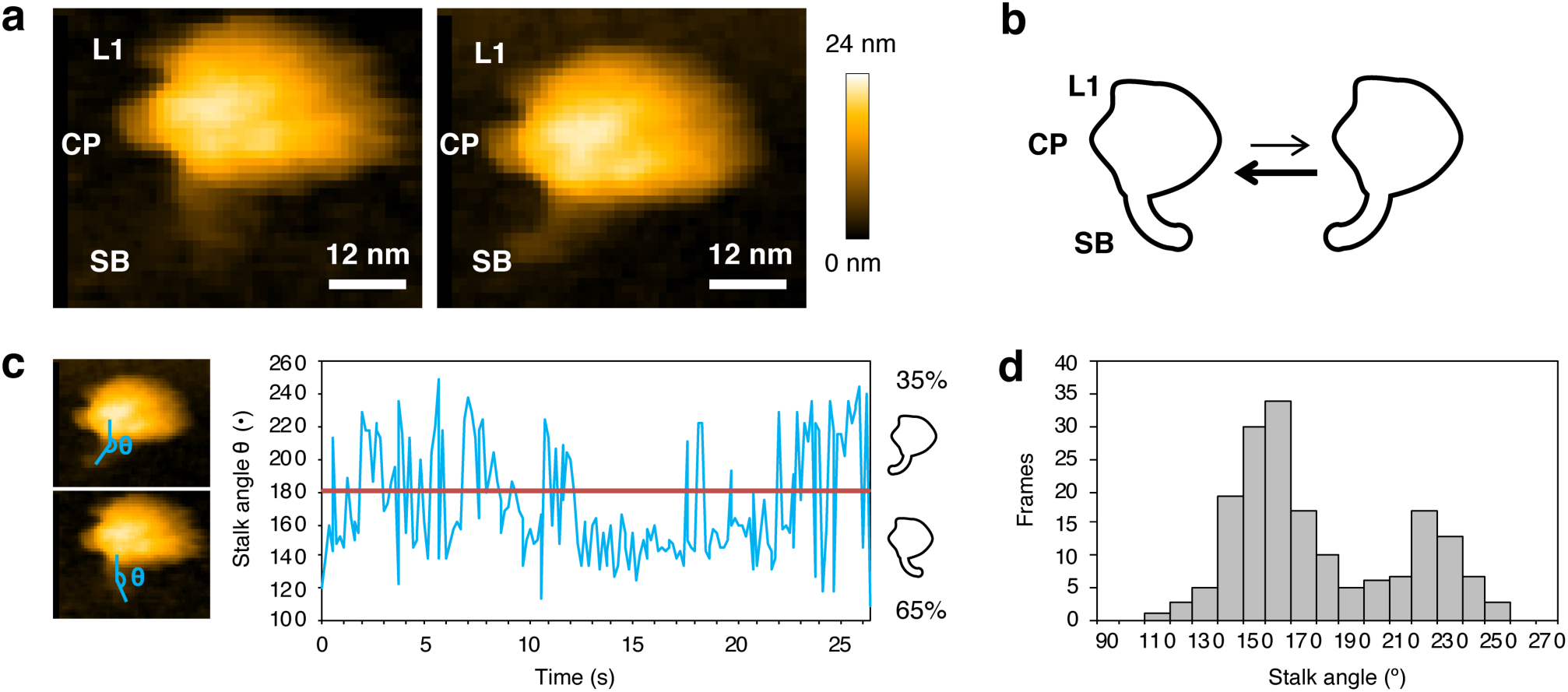
Structural dynamics of the ribosomal P-stalk base on the archaeal 50S subunit. **a**, HS-AFM images of the ribosomal stalk base in the canonical (left) and flipped (right) states (L1: L1 stalk, CP: central protuberance, SB: P-stalk base). Scan area, 60 × 50 nm with 50 × 40 pixels; scan speed, 150 ms/frame; scale bar, 12 nm. **b**, Schematic representations of two conformational states of the ribosomal P-stalk base, related to **a. c**, Time course of the structural rearrangements of the ribosomal P-stalk, related to **a** and Supplementary movie 2. The angle between the proximal and distal regions of the stalk, θ (°) are indicated by the cyan-line (θ) in left two images. **d**, Histogram of the stalk base angles (°), related to **c**.

To confirm whether the protrusion observed in Fig. 1a is the ribosomal P-stalk, we used a hybrid ribosome system that allows removing and reconstitution of the ribosomal stalk *in vitro* (31). In this system, the bacterial stalk proteins (bL10 and bL12) are removed from the *E. coli* 50S subunit lacking the bacterial ribosomal protein L11 (bL11). The resulting *E. coli* 50S core can be reconstituted by any type of stalk complex and L11. We previously reconstituted the hybrid 50S subunit, in which the archaeal ribosomal P-stalk complexes and archaeal L11 (aL11) are introduced into the *E. coli* 50S core and demonstrated that the hybrid 70S ribosome exhibits translation elongation depending on archaeal trGTPases (13, 32, 33). Here, according to the previous report, we prepared the *E. coli* 50S core and reconstituted the hybrid 50S subunit by adding archaeal aP0•(aP1•aP1)_3_ P-stalk complex and aL11. HS-AFM observation of the 50S core alone showed a spherical shape without protrusions (Fig. 1c). On the other hand, similar to Fig. 1a, we succeeded in detecting the protrusions in HS-AFM images of the hybrid 50S subunit (Fig. 1d). These protrusion structures were detected in about 80% of the total hybrid 50S subunits observed (Fig. 1e, Supplementary Fig. 3). These results reveal that the protrusion observed in Fig. 1a is the ribosomal P-stalk base (see Fig. 1a, left), and that HS-AFM is useful for the detection and investigation of the ribosomal P-stalk on the 50S ribosomal subunit.

**Figure 3.**
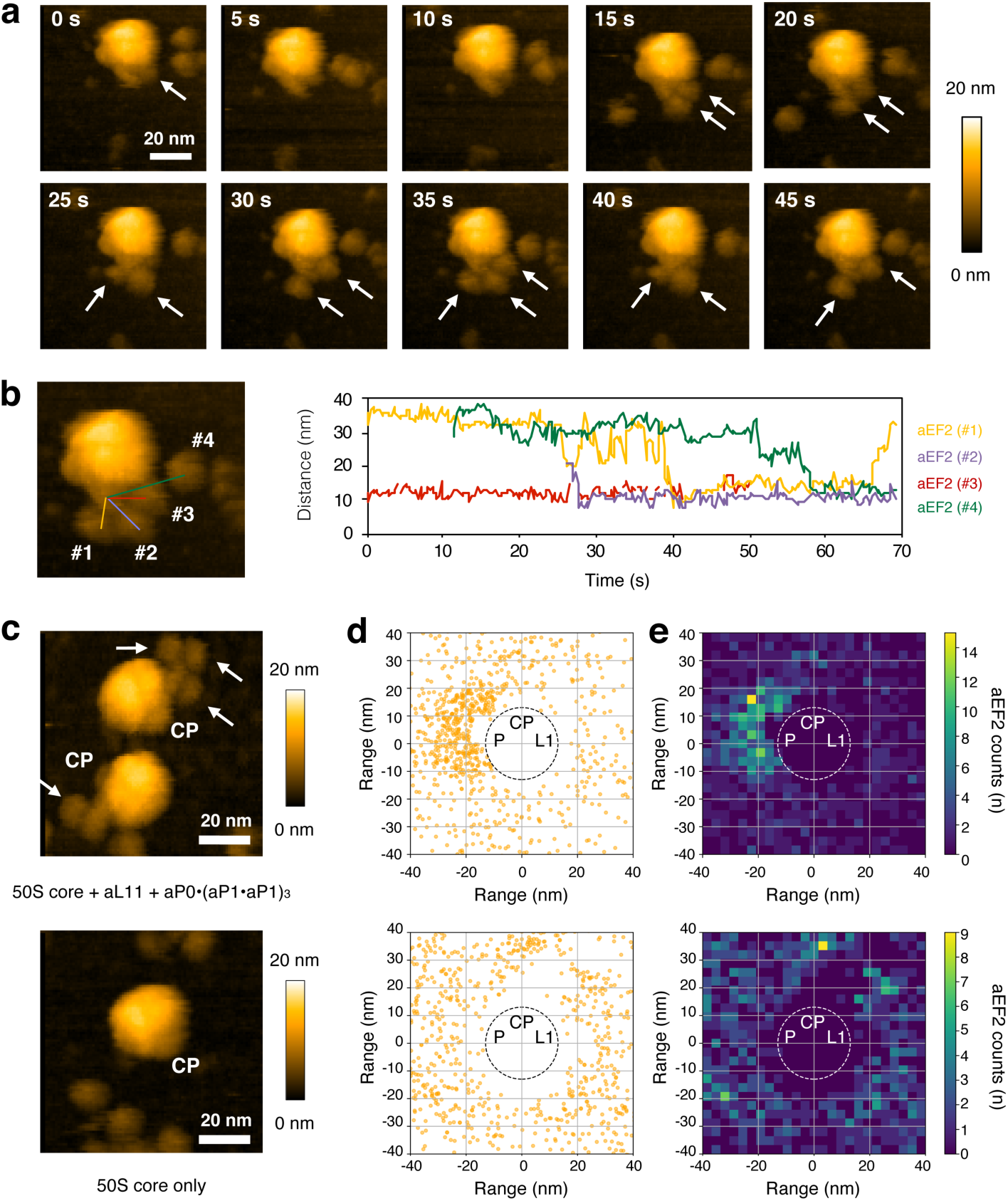
The ribosomal stalk simultaneously recruits multiple aEF2•GTP to the ribosome to form the factor pool. **a**, Sequential HS-AFM images of the hybrid 50S subunit after the addition of aEF2•GTP on the AP-mica surface. aEF2•GTP molecules bound to the ribosomal stalk are indicated by white arrows. Scan area, 80 × 80 nm with 90 × 90 pixels; scan speed, 250 ms/frame; scale bar, 20 nm. **b**, A HS-AFM image related to Fig. 3a (left). Time courses of the distance between the ribosomal P-stalk and individual aEF2•GTP molecules (#1: orange, #2: purple, #3: red, #4: green) related to **a** (right). **c**, Typical HS-AFM images of the hybrid 50S (above) or 50S core (below) in the presence of 50 nM aEF2•GTP. Scan area, 80 × 80 nm with 90 × 90 pixels; scan speed, 250 ms/frame; scale bar, 20 nm. aEF2 molecules bound to the ribosomal stalk are indicated by white arrows. **d, e**, dot plots (**d**) and heat maps (**e**) of aEF2 distribution measured from the center of the hybrid 50S (930 aEF2 molecules and 156 subunits were analyzed) or 50S core (1,316 aEF2 molecules and 112 subunits were analyzed), related to **c**. Dotted circles approximately represent the 50S subunit (L1: L1 stalk, CP: central protuberance, P: P-stalk base).

### Structural dynamics of the ribosomal P-stalk on the large subunit

Previous structural studies have shown only part of the ribosomal stalk in the ribosome-bound states. To date, three-dimensional models of the ribosome containing the ribosomal stalk have been generated by superimposing the crystal structure of the isolated ribosomal stalk onto the ribosome (4, 9). Therefore, our knowledge about the structural dynamics of the ribosomal stalk in the ribosome-bound states is limited. To investigate the structural dynamics of the ribosomal P-stalk on the large subunit, we analyzed HS-AFM images of the archaeal 50S subunit. First, we confirmed that the archaeal ribosomal P-stalk has a canonical state in agreement with previous structural models, in which the ribosomal stalk binds to the 23S rRNA through the N-terminal domain of aP0, and the C-terminal side of aP0 is oriented away from the CP (Fig. 1a; left, Fig. 2a). Intriguingly, an unexpected new state was observed in which the ribosomal P-stalk tip flips toward the CP (Fig. 2a; right). In the HS-AFM movie, the ribosomal P-stalk was moving back and forth between the canonical and flipped states with a time ratio of 65% and 35%, respectively (Fig. 2b, c, Supplementary Movie 2). The histogram of the angle between the proximal and distal regions of the P-stalk from each HS-AFM image showed two peaks, indicating that the ribosomal P-stalk base has at least two conformational states, the predicted canonical state and the newly observed flipped state (Fig. 2d). These results showed that the ribosomal P-stalk has a flexible property not only at the C-terminal disordered regions but also nearer to the stalk base.

### aEF2 assembles to the ribosomal P-stalk on the large subunit

Although the factor-pooling mechanism is hypothesized as one of the functions of the ribosomal stalk, conclusive evidence in support of this mechanism has not been reported. To address this, visualization of the distribution of trGTPases around the ribosome is required at a resolution of nanometer-order. HS-AFM can directly visualize the structural dynamics of proteins, including the surrounding environment with nanometer-order resolution (34–39). We here tested whether the ribosomal P-stalk participates in the formation of a trGTPases factor pool around the ribosome. To this end, we used the hybrid 50S subunit system, which can easily detect the binding of archaeal trGTPases to the ribosomal P-stalk bound to the large ribosomal subunit (13) (Supplementary Fig. 4). First, the hybrid 50S subunit was immobilized on the AP-mica surface in the presence of GTP, and aEF2 was added to the sample chamber at 50 nM to give a 10:1 stoichiometry with the ribosome during the HS-AFM observation. After the addition of aEF2, molecules with a size equivalent to aEF2 (∼7 nm) appeared on the AP-mica surface (Supplementary Fig. 5a). HS-AFM analysis showed that multiple aEF2 molecules gradually assembled to the hybrid 50S subunit (Fig. 3a, b, Supplementary Fig. 5b, Supplementary Movie 3, 4). Quantification of aEF2 distribution around the subunit particles revealed that aEF2 molecules preferentially distributed near the ribosomal P-stalk 6.6 fold relative to the opposite control area (Fig. 3c–e upper panels, Supplementary Fig. 5c). The preferential aEF2 distribution was also observed in the presence of GDP, in which the aEF2 distributed near the ribosomal P-stalk approximately 4.3 fold relative to the opposite control area (Supplementary Fig. 5d). Measuring the distances from the centers of recruited aEF2 molecules to the center of the P-stalk revealed a mean distance of 12.9 nm for aEF2•GTP and 11.5 nm for aEF2•GDP, respectively (Supplementary Fig. 5f, g). To investigate whether the preferential aEF2 distribution was dependent on the ribosomal P-stalk, we performed HS-AFM observation using the *E. coli* 50S core instead of the hybrid 50S subunit. As a result, an unbiased, even distribution of aEF2 was observed (Fig. 3c–e lower panel, Supplementary Fig. 5e, h, i). These results demonstrate that the ribosomal P-stalk recruits multiple aEF2 molecules around itself for increasing the local concentration of aEF2 in a manner consistent with the factor-pooling hypothesis.

**Figure 4.**
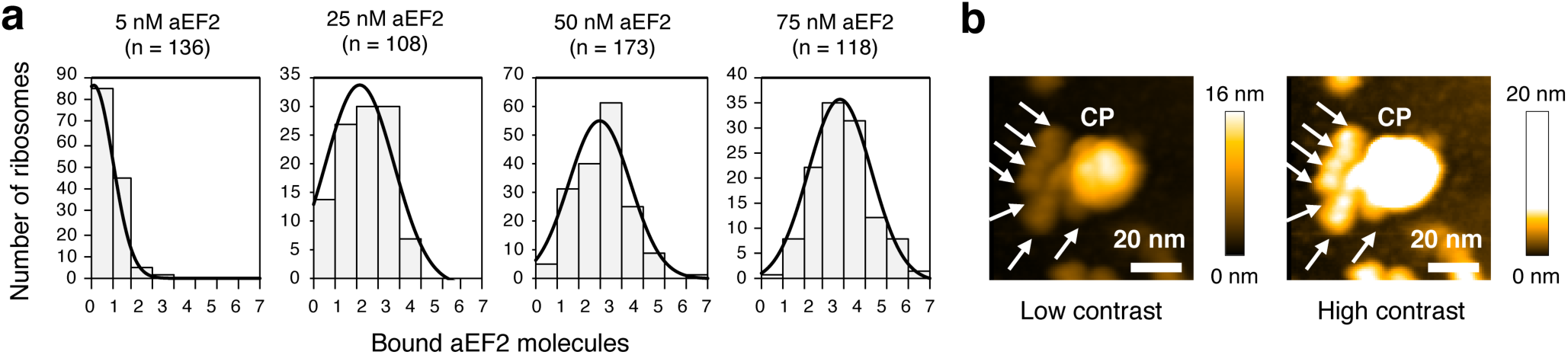
The ribosomal stalk simultaneously binds to multiple aEF2 molecules in a concentration dependent manner. **a**, Histograms of the binding numbers of aEF2 to the ribosomal stalk. The hybrid 50S subunits (5 nM) were incubated with various concentrations of aEF2•GTP (5, 25, 50, and 75 nM) and immobilized to the AP-mica surface. After washing, the number of aEF2 molecules bound to each ribosomal stalk was counted by HS-AFM (n: number of 50S particles counted). **b**, HS-AFM images of the hybrid 50S bound to seven aEF2 molecules with low (left) and high (right) contrasts. Scan area, 80 × 80 nm with 90 × 90 pixels; scan speed, 250 ms/frame; scale bar, 20 nm. aEF2 molecules bound to the ribosomal stalk are indicated by white arrows.

**Figure 5.**
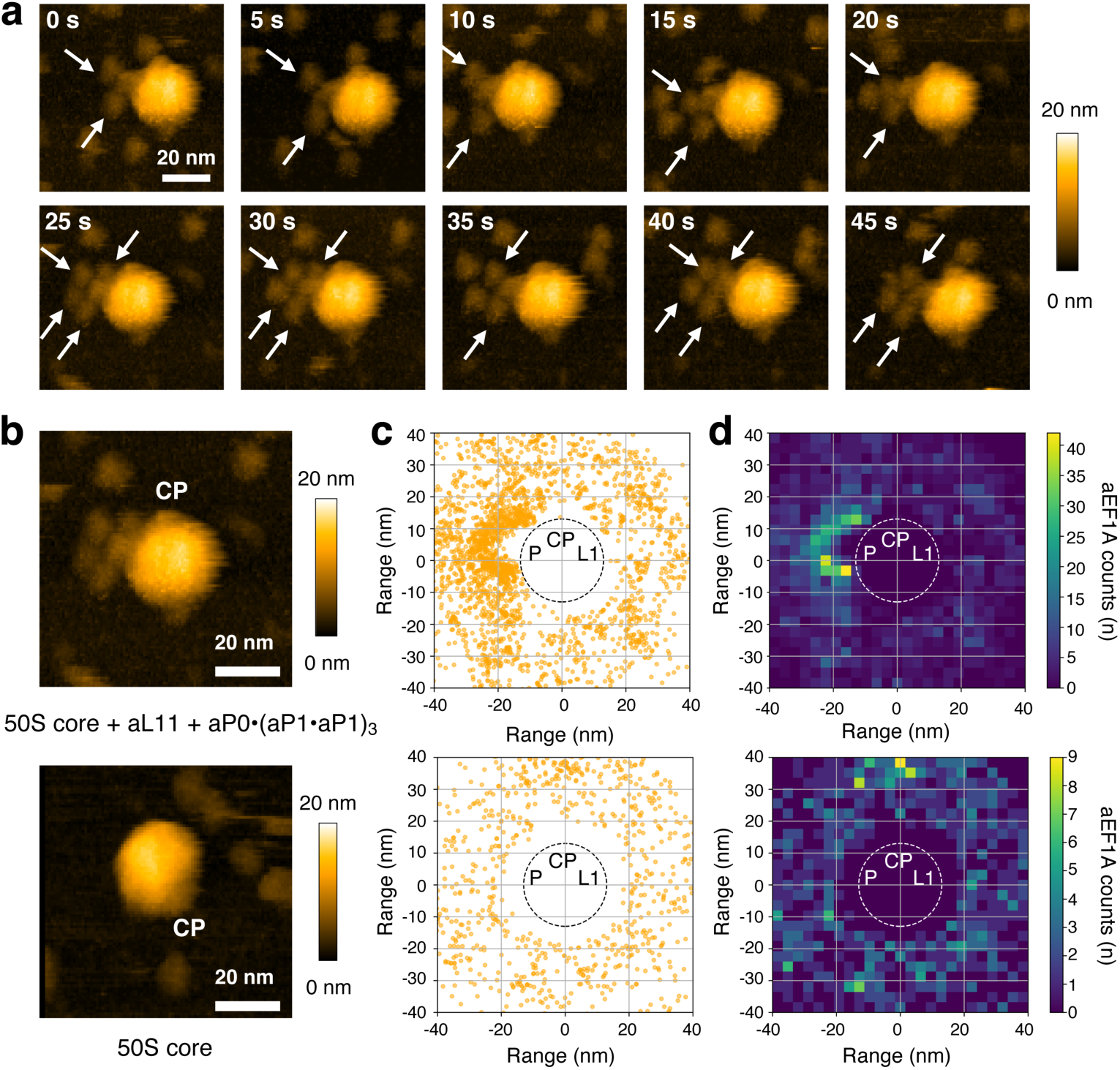
Preferential distribution of aEF1A•GTP around the 50S subunit dependent on ribosomal stalk. **a**, Sequential HS-AFM images of the hybrid 50S subunit in the presence of 50 nM aEF1A•GTP on the AP-mica surface. aEF1A•GTP molecules bound to the ribosomal stalk are indicated by white allows. Scan area, 80 × 80 nm with 90 × 90 pixels; scan speed, 250 ms/frame; scale bar, 20 nm. **b**, Typical HS-AFM images of the hybrid 50S (above) or 50S core (below) in the presence of 50 nM aEF1A•GTP. Scan area, 80 × 80 nm with 90 × 90 pixels; scan speed, 250 ms/frame; scale bar, 20 nm. aEF1A•GTP molecules bound to the ribosomal stalk are indicated by white arrows. **c, d**, dot plots (**c**) and heat maps (**d**) of aEF1A•GTP distribution around the hybrid 50S (2,490 aEF1A molecules and 278 subunits were analyzed) or 50S core (937 aEF1A molecules and 141 subunits were analyzed), related to **c**. Dot circles approximately represent the 50S subunit (L1: L1 stalk, CP: central protuberance, P: P-stalk base).

### The ribosomal P-stalk simultaneously binds to multiple aEF2 molecules

The archaeal ribosomal P-stalk has seven copies of the C-terminal region, which presumably bind to trGTPase individually (8, 9). Nomura et al. previously observed that the isolated archaeal ribosomal P-stalk simultaneously binds to at least three aEF2 molecules by ultracentrifugation experiments (13). However, the maximum number of aEF2 molecules that can be simultaneously bound to the ribosomal P-stalk is unclear. To examine this question, we pre-incubated hybrid 50S subunits with various concentrations of aEF2 in the presence of GTP and immobilized them on AP-mica. After washing, aEF2 molecules bound to the ribosomal P-stalk were counted. Unfortunately, when the aEF2 concentration was higher than 80 nM, it became difficult to distinguish between aEF2 bound to the ribosomal P-stalk and free aEF2 on the AP-mica during HS-AFM observation. Therefore, we observed in the range of 5–75 nM of aEF2, and we counted more than 100 50S particles at each concentration. The number of aEF2 molecules bound around to the ribosomal P-stalk increased in a concentration-dependent manner, and up to seven aEF2 molecules simultaneously bound to the ribosomal P-stalk (Fig. 4a, b). The results reveal that each P-stalk C-terminal region can independently recruit an aEF2 molecule, and the number of recruited aEF2 molecules depends on the concentration of aEF2.

### aEF1A assembles to the ribosomal P-stalk both in GTP-and GDP-bound forms

Efficient binding and dissociation of EF1A•GTP•aa-tRNA ternary complex to the ribosome are critical for correct codon reading at the A-site and for improving translation accuracy. In previous studies, we have demonstrated that the ribosomal P-stalk is also involved in the recruitment of EF1A•GTP•aa-tRNA and the activation of GTP hydrolysis (33, 40). We next visualized the binding of aEF1A to the ribosomal P-stalk on the large subunit. Using HS-AFM, we observed that aEF1A (∼5 nm) transiently interacts with the ribosomal P-stalk on the 50S subunit in the presence of GTP (Fig. 5a, Supplementary Fig. 6, Supplementary Movies 5). In this condition, aEF1A molecules associated near the ribosomal P-stalk 4.2 fold more often relative to the opposite control area, forming a pool of aEF1A (Fig. 5b–d upper panel, Supplementary Fig. 6c), and the histogram of distances from the center of each aEF1A molecule to the center of the P-stalk showed a mean distance of 9.5 nm for the P-stalk-bound aEF1A (Supplementary Fig. 6d). The preferential distribution of aEF1A•GTP was not observed using the 50S core (Fig. 5b–d lower panel, Supplementary Fig. 6e, f). Therefore, similar to aEF2, aEF1A localizes near the factor-binding center in a ribosomal P-stalk dependent manner. The binding and dissociation of aEF1A•GTP to the ribosomal P-stalk (Supplementary Movies 5, 6) seems to take place more rapidly than aEF2•GTP (Supplementary Movie 4). The short binding lifetime of aEF1A•GTP to the ribosomal P-stalk may reflect the necessity for kinetically rapid association and dissociation to allow efficient codon-anticodon reading tests.

**Figure 6.**
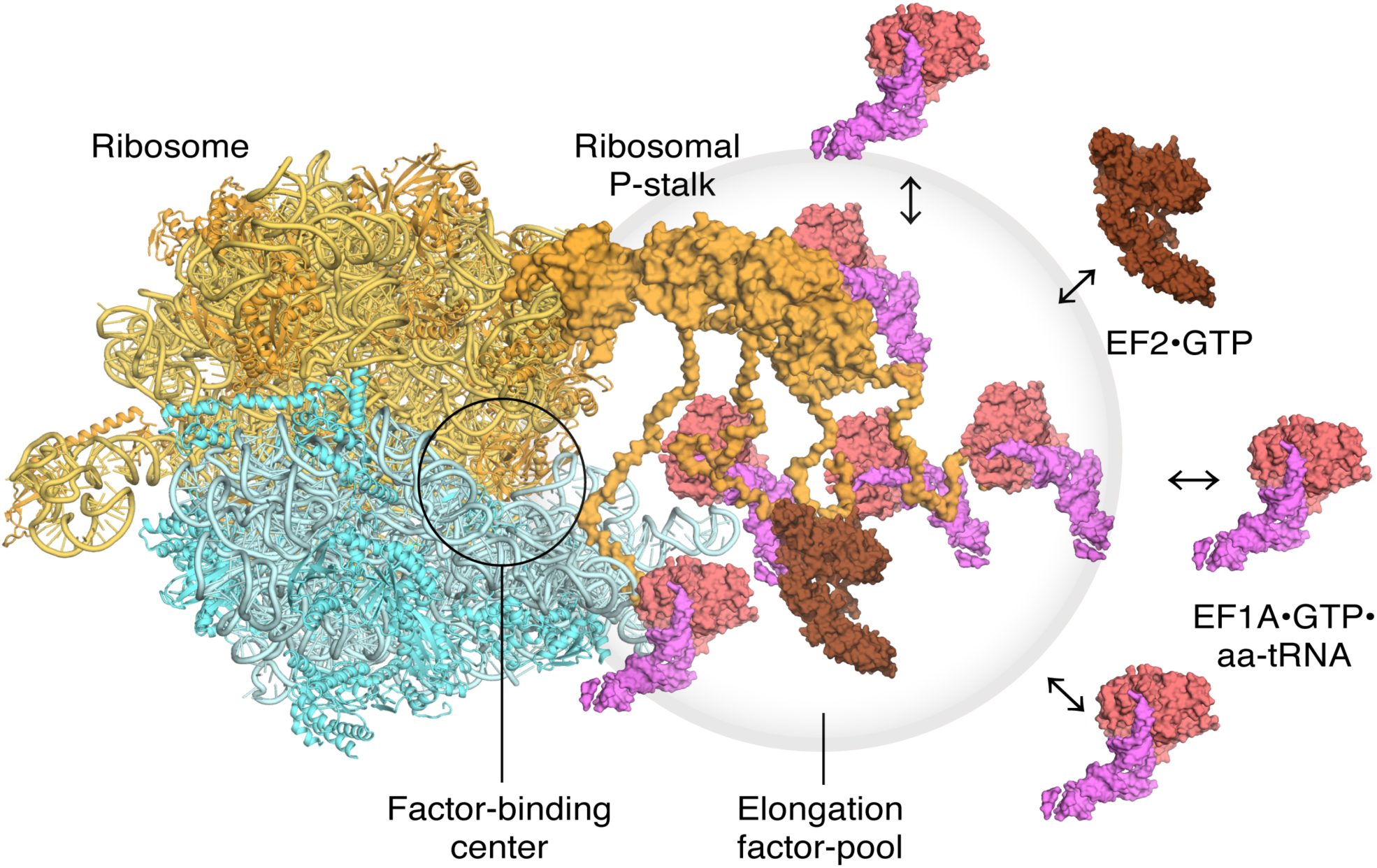
Model of translating ribosomes and elongation factors. EF1A•GTP•aa-tRNA and EF2 assemble to the ribosomal stalk on the translating ribosome. The translation factor pool contributes to efficient protein synthesis in a crowded intracellular environment.

It has been demonstrated that the conformation of the GDP-bound form of aEF1A and its binding mode to the C-terminal region of the aP1 stalk are somewhat different from those of the GTP-bound form (40, 41). We thus analyzed the binding of aEF1A•GDP to the ribosomal P-stalk on the large subunit. The results also showed the preferential distribution of aEF1A•GDP in proximity to the P-stalk, revealing that both GTP-and GDP-binding forms of aEF1A localize around the factor-binding center of the ribosome (Supplementary Fig. 7a). The aEF1A•GDP associated near the ribosomal P-stalk 2.6 fold more often relative to the opposite control area (Supplementary Fig. 7b, c). These GTP-and GDP-dependent factor-pool formations are consistent with a similar binding affinity of aP1 to aEF1A•GTP and aEF1A•GDP (40, 41) (Supplementary Table 1). Since the formation of the aEF1A-pooling was not observed in nucleotide-free conditions, the present HS-AFM data show that aEF1A assembly is dependent on GTP and GDP (Supplementary Fig. 7d–g). Taken together, these results indicate that the ribosomal P-stalk increases the local concentration of aEF1A near the factor-binding center, in agreement with the factor-pooling hypothesis to efficient decoding by EF1A•GTP•aa-tRNA and translation elongation.

## Discussion

The ribosomal stalk plays an essential role in the efficient binding of trGTPases to the ribosome, but its mechanism of action is not fully understood. In this study, using HS-AFM, we directly visualized (i) the structural dynamics of the archaeal ribosomal P-stalk on the 50S subunit, (ii) the assembly processes of trGTPases to the ribosomal P-stalk, and (iii) the P-stalk-dependent preferential distribution of trGTPases around the factor-binding center. These results reveal that the ribosomal P-stalk forms a trGTPase pool near the factor-binding center for efficient trGTPases turnovers. It must be noted that the trGTPases factor pool observed in this study was formed on the hybrid 50S subunit immobilized on AP-mica, therefore we did not directly visualize the full biological translation processes mediated by the 70S ribosome and other cofactors. However, the 30S subunit does not influence the interactions between the ribosomal P-stalk and trGTPases, and the hybrid system we used accurately recapitulates the functional role of the archaeal and eukaryotic ribosomal P-stalk in translation elongation (10, 31–33, 40). Therefore, we believe that the formation of the trGTPase factor pool we observed here presumably reflects the actual translation elongation status is likely to occur during actual biological translation elongation.

Using HS-AFM, we first observed an unexpected structural flexibility of the ribosomal P-stalk base on the ribosome (Fig. 2). This structural rearrangement of the P-stalk base has not been detected in X-ray crystallography and cryo-EM studies of ribosomal particles. Our present study demonstrates that the structural rearrangements of the ribosomal P-stalk base take place at a high frequency that we hypothesize to play some functional role. A reasonable hypothesis is that the movement of the P-stalk base expands the range of movement of the C-terminal part of each stalk protein, resulting in increased frequency of the stalk interaction with translation factors around the ribosome (14).

HS-AFM directly showed aEF2•GTP and aEF1A•GTP preferentially distributed near the ribosomal P-stalk. These increases in the local concentration of aEF2/aEF1A depended on the presence of the ribosomal P-stalk, suggesting that the P-stalk works as the factor-pooling platform. We hypothesize that this mechanism is driven by two key properties of the P-stalk: the rapid flipping movement we described above, and the presence of multiple copies of the flexible C-terminal region for parallel binding of trGTPases. HS-AFM analysis showed the mean distances from the center of the P-stalk base to the center of each trGTPase were 12.9 nm for aEF2•GTP and 9.5 nm for aEF1A•GTP, respectively (Supplementary Fig. 5f, 6d). These observations are consistent with the fact that the C-terminal regions of eukaryotic P1/P2, which is homologous to aP1, can extend up to ∼12.5 nm away from its N-terminal dimerization domain (14). The number of C-terminal regions of the ribosomal stalk is correlated to the trGTPase activity and polypeptide elongation rate *in vitro* (9, 13, 33, 42). In addition, Wawiórka et al. reported that decreasing the number of C-terminal regions of the ribosomal stalk in *Saccharomyces cerevisiae* reduces translation accuracy (43). From these lines of evidence, we infer that the ribosomal stalk participates in forming a pool of different EF1A•GTP•aa-tRNA ternary complexes around the P-stalk for efficient codon-anticodon reading tests, and that maintaining this aEF1A/aEF2 pool is important for rapid and accurate translation elongation in crowded intracellular environments (Fig. 6).

Recently, using single-molecule fluorescence method, Mustafi et al. tracked a single molecule of EF-Tu was fused with the fluorescent protein mEos2 in *E. coli* cells, with resolution on the micrometer order, and classified EF-Tu into two groups based on differences in their diffusion rates (44). In their observations, EF-Tu with a slow diffusion rate (approximately 60% of the total EF-Tu) was co-localized with the ribosome in a manner dependent on interaction with the L7/L12-stalk, and the average stoichiometry of EF-Tu binding to the ribosome was estimated to be 3.5 molecules of EF-Tu per ribosome. Although it is unknown whether EF-Tu has a preferential distribution around the factor-binding center based on these experiments, their study is consistent with our present observations and suggests that the factor-pooling mechanism visualized by HS-AFM is biologically relevant.

In translation initiation, the ribosomal stalk interacts with the GTPase initiation factor IF2 (aIF5B in archaea and eIF5B in eukaryotes) to promote large and small subunit joining and selectivity for the initiation AUG codon (45, 46). The ribosomal stalk also plays a functional role in the recruitment of ribosome recycling factor ABCE1 and its ATP hydrolysis in archaea and eukaryotes (47). Point mutations disrupting the binding of the ribosomal stalk to eIF5B and ABCE1 inhibit cell viability, suggesting that the ribosomal stalk maintains these functional interactions with translation initiation and recycling factors *in vivo* (46, 47). Therefore, at least four translation factors appear to compete with each other as the binding partners for the ribosomal stalk. In previous *in vitro* studies, it has been found that the binding affinity of the C-terminal region of the stalk to each translation factor does not differ significantly (Supplementary Table S1) (41, 46–48). Given that the number of translation factors bound around the ribosomal stalk is concentration-dependent (Fig. 4), it is likely that the identities and numbers of factors that bind to the multiple stalk C-terminal regions depend on the local concentration of each factor. Supporting this notion is the fact that EF1A, which is the most likely trGTPase to interact with the ribosome in translation, is one of the most abundant proteins in the cell.

Unfortunately, our HS-AFM analyses do not provide any information about the action of the ribosomal stalk and trGTPases once bound to the factor-binding center of the ribosome. Wang et al., using an *E. coli* system, reported that the stalk may have the ability to stabilize the GTP-bound trGTPases on the factor-binding center via interaction of the stalk with ribosomal protein uL11, possibly promoting trGTPases-dependent polypeptide synthesis and/or GTPase activation (49). Mohr et al. showed that the stalk participates in the release of inorganic phosphate from EF-G after GTP hydrolysis (3). In the present HS-AFM study, we clearly show that each copy of aP0 and aP1 also has the ability to interact with aEF1A•GDP or aEF2•GDP molecules, suggesting that the ribosomal P-stalk keeps its aEF1A/aEF2 pool bound nearby even after their dissociation from the ribosomal factor-binding center after GTP hydrolysis. The retention of the aEF1A•GDP/aEF2•GDP pool may also allow rapid re-use of trGTPases without having to dissociate and re-associate for the nucleotide exchange reaction. Therefore it seems likely that, by making and maintaining the aEF1A/aEF2-pool, the stalk efficiently promotes not only the recruitment of GTP-bound trGTPases to the factor-binding center and their actions including GTP hydrolysis on the ribosome, but also the retention of GDP-bound trGTPases after their dissociation from the ribosomal factor-binding center.

In conclusion, this study provides the first visual evidence for the trGTPase factor-pooling mechanism by the ribosomal P-stalk. Translation is a collection of dynamic processes in which the ribosome undergoes sequential conformational changes and interacts with many accessory factors. Since HS-AFM has advantages in observation of not only structural dynamics but also spatial distributions of biomolecules, future work with HS-AFM will provide further important information to understand the dynamic behaviors of these complex translational machineries.

## Materials and Methods

### Sample preparation

The plasmids for expression of *Pyrococcus horikoshii* aP0, aP1, aL11 were constructed, and the proteins were expressed and purified as described previously (9, 32). The plasmids for expression of *Pyrococcus furiosus* aEF2 and aEF1A were constructed, and the proteins were expressed using an *Escherichia coli* expression system and isolated as described previously (33, 40). The prepared aEF1A sample contained tightly bound GDP (40). In order to observe the dynamics of GTP-bound aEF1A binding to the ribosomes, we prepared a nucleotide-free aEF1A (NF) by bacterial alkaline phosphatase treatment (50). The 50S ribosomal subunits from *Pyrococcus furiosus* were prepared as described previously (13). The 50S and 30S ribosomal subunits from *E. coli* Q13 strain and the 50S subunit lacking bL11 from *E. coli* AM68 strain were prepared as described previously (31, 51). The *E. coli* 50S core were prepared as described previously (31). The hybrid 50S subunits were reconstituted by mixing the *E. coli* 50S core, with purified archaeal aL11 protein and purified aP0•(aP1•aP1)_3_ complexes as described previously (32). In brief, 10 pmol of 50S cores were incubated with 20 pmol of aL11 and 20 pmol of aP0•(aP1•aP1)_3_ complex in 10 *µ*l of buffer A (20 mM Tris-HCl, pH 7.6, 10 mM MgCl2, 50 mM NH4Cl) for 10 min at 37°C.

### Native-PAGE

5 pmol of the 50S cores or hybrid 50S subunits were incubated without or with 5, 10, 15, 20, 25, 30, 35, 40, 45 pmol of aEF2 in 10 *µ*l of buffer A for 10 min at 37°C. After incubation, 0.7 *µ*l of loading buffer with dye [50% (v/v) glycerol, 0.1% (w/v) bromophenol blue, 0.1% (w/v) xylene cyanol] was added. The total sample solution was subjected to acrylamide-agarose composite gel electrophoresis, as described previously (10).

### HS-AFM observations

A laboratory-built tapping mode high-speed AFM was used (52, 53). The short cantilevers used (BL-AC10DS-A2) were purchased from Olympus. The spring constant of the cantilever was ∼100 pN/nm. The resonant frequency and the quality factor of the cantilever in liquid were ∼500 kHz and ∼1.5, respectively. An amorphous carbon tip was fabricated on the original AFM tip by electron beam deposition (EBD). The length of the additional AFM tip was ∼500 nm, and the radius of the apex of the tip was ∼4 nm. The free oscillation amplitude of the cantilever was 2 nm, and the set-point amplitude was set to 90% of the free amplitude. For HS-AFM observations of the 50S subunits, a mica surface was treated for 5 min with 0.01% (v/v) (3-aminopropyl)triethoxysilane (APTES) (Sigma-Aldrich). HS-AFM observations of the 50S subunits from *Pyrococcus furiosus* or pre-assembled hybrid 50S subunits were performed as follows. The ribosome mixture containing *P. furiosus* 50S (5 nM) or the hybrid 50S (5 nM) subunits was incubated in buffer A for 10 min at 37°C. 2 *µ*l of the ribosome mixture was placed on the AP-mica and incubated for 5 min at room temperature. After washing with 20 *µ*l of buffer A, the stage was immersed in 60 *µ*l of buffer A filled in the liquid cell of the cantilever holder and HS-AFM observation was started.

Observation of aEF2 binding to hybrid 50S subunits was performed as follows. While observing the hybrid 50S subunits under the conditions described above, aEF2 and GTP were added to the liquid cell at final concentrations of 50 nM and 1 mM, respectively. After adding aEF2 and GTP, HS-AFM observation was started immediately.

Counting of the ribosomal stalk bound aEF2 was performed as follows. The mixture containing the hybrid 50S subunits (5 nM) and aEF2 (5, 25, 50, and 75 nM) was incubated in buffer A containing 1 mM GTP for 10 min at 37 °C. 2 *µ*l of this ribosome mixture was placed on the AP-mica and incubated for 5 min at room temperature. After washing with 20 *µ*l of buffer A, the stage was immersed in 60 *µ*l of buffer A in the liquid cell of the cantilever holder and HS-AFM observation was started.

The observation of aEF1A was performed as follows. The mixture containing the hybrid 50S subunits (5 nM) and aEF1A (50 nM) was incubated in buffer A in the presence or absence of 1 mM GTP or GDP for 10 min at 37 °C. 2 *µ*l of this ribosome mixture was placed on the AP-mica and incubated for 5 min at room temperature. After washing with 20 *µ*l of buffer A, the stage was immersed in 60 *µ*l of buffer A and HS-AFM observation was started.

### Analyzing of HS-AFM images

HS-AFM images were viewed and analyzed using a laboratory-built software, Kodec4.5.7.39 (54). A flattening filter to make the XY-plane flat was applied to individual images. The heights of ribosomal subunits were determined using a cross-sectional analysis. Angles of the ribosomal stalk base were determined as follows. First, XY coordinates of three points (root, bend, and the tip of the ribosomal stalk) were obtained from individual images. The stalk base angles were calculated from these three points.

The distributions of aEF2 and aEF1A were quantified as follows. First, the XY coordinates of the center of the ribosome (CR), the central protuberance (CP), and the center of the translational factors (CT) were manually obtained from individual HS-AFM images. For each frame, all coordinates were translated so that the CR was placed on the origin, and all coordinates were rotated so that the CP was placed on the positive Y axis. Then, the CT values from all images were plotted as XY coordinates which were used to create a heatmap showing the distribution of the translation factors.

### Data availability

The data sets generated during the current study are available from the corresponding author upon request.

## Acknowledgment

We thank Dr. Steven J. McArthur for critical reading and the English language improvement of the manuscript. We thank Drs. Yoshizumi Ishino and Sonoko Ishino for kindly providing the culture pellets of *Pyrococcus furiosus*. We thank the technical support staff for the HS-AFM, including Ms. Aimi Makino, Drs. Toshio Ando, and Takayuki Uchihashi. This work was supported by JSPS (KAKENHI 20J00036 to H.I., 19H03155 to T.U., 18H05269 to N.K.) and JST-CREST (JPMJCR1762 to N.K.). This work was supported in part by Bio-SPMs collaborative research of WPI-NanoLSI, Kanazawa University.

## Author contributions

All authors designed experiments. H.I. performed all experiments and statistical analysis of the data. All authors edited the manuscript.

