## Supplemental Information for "Direct visualization of translational GTPase factor-pool formed around the archaeal ribosomal P-stalk by high-speed atomic force microscopy"

**Supplementary Fig. 1. HS-AFM images of the *Escherichia coli* 50S and 30S ribosomal subunits on AP-mica surface. a,** Wide-range HS-AFM images of the 50S subunits from *E. coli* on AP-mica surface. Scan area, 160 x 160 nm with 100 x 100 pixels; scan speed, 330 ms/frame; scale bar, 40 nm. **b, c,** Typical HS-AFM images of the 50S and 30S subunits from *E. coli* on AP-mica surface (L1: L1 stalk, CP: central protuberance, SB: stalk base). Scan area, 80 x 80 nm with 80 x 80 pixels; scan speed, 250 ms/frame; scale bar, 20 nm. **d, e,** High-contrasted HS-AFM images, related to **b** and **c**. **f, g,** Histograms of the heights of individual 50S (**f**) and 30S (**g**) subunits in the HS-AFM images. **h, i,** Crystal structures of the 50S (h: orange) and 30S (i: blue) from *E. coli* [PDB ID: 4YBB] and its expected orientation on the AP-mica surface.

**Supplementary Fig. 2. Comparison of the ribosomal stalk between bacteria and archaea. a,** Sequence alignment of the ribosomal proteins bL10 and aP0. The sequences of bL10 from *Escherichia coli* (*Eco*) and *Thermotoga maritima* (*Tma*), and aP0 from *Pyrococcus horikoshii* (*Pho*) were compared by CLUSTALW (https://www.genome.jp/tools-bin/clustalw). **b,** Structural model of the *E. coli* bL10•(bL12•bL12)$_2$ L7/L12-stalk complex, based on the *Thermotoga maritima* bL10•(bL12•bL12)$_3$ L7/L12-stalk complex [PDB ID: 1ZAX]. **c,** Structure of the archaeal aP0•(aP1•aP1)$_3$ P-stalk complex from *Pyrococcus horikoshii* [PDB ID: 3A1Y]. In **b** and **c**, the NTD of bL10/aP0 proteins are colored blue. The bL12- and aP1-binding helix of bL10/aP1 are colored cyan. bL12 and aP1 dimer 1, 2, and 3 are colored yellow, orange, and red, respectively. N and C indicate the N- and C-terminus of bL10/aP0, respectively. The C-terminus of the ribosomal stalk are not shown.

**Supplementary Fig. 3. Typical HS-AFM images of the 50S cores and the hybrid 50S subunits, related to figures 1c and 1d, respectively.** The 50S core were incubated without (a) or with (b) ribosomal protein aL11 and aP0•(aP1•aP1)$_3$ complex and immobilized on the AP-mica surface. Scan area, 80 x 80 nm with 80 x 80 pixels; scan speed, 250 ms/frame; scale bar, 20 nm. White arrows indicate the ribosomal P-stalk base.

**Supplementary Fig. 4. Interaction of the hybrid 50S subunit with aEF2.** The hybrid 50S subunit (5 pmol) was incubated with aEF2 (5-45 pmol) in the absence (lane 1) or presence (lanes 2–11) of aL11 and aP0•(aP1•aP1)$_3$ and subjected to Native-PAGE. The gel was stained by AzurB.

**Supplementary Fig. 5. HS-AFM images of the hybrid 50S subunits in the presence of aEF2, related to Fig. 3. a,** Histogram of heights of aEF2 molecules. **b,** High-contrast sequential HS-AFM images of the hybrid 50S subunit after the addition of aEF2 on the AP-mica surface, related to Fig. 3a. aEF2 molecules bound to the ribosomal stalk are indicated by white arrows. Scan area, 80 x 80 nm with 90 x 90 pixels; scan speed, 250 ms/frame; scale bar, 20 nm. **c–e,** Dot plots (left) and heat maps (right) of aEF2•GTP molecules around the hybrid 50S subunit (**c**), aEF2•GDP molecules around the hybrid 50S subunit (**d**), and aEF2•GTP molecules around the 50S core (**e**), related to Fig. 3d. The area near the P-stalk side (PS; radius, 20 nm) or the area opposite the P-stalk (OP; radius, 20 nm) are represented by red and blue circles, respectively. The ratios of the number of molecules of aEF2 (PS/OP) are 6.6 (**c**), 4.3 (**d**), and 1.0 (**e**), respectively. In **d**, 1,632 aEF2 molecules and 245 subunits were analyzed. **f–h,** Histograms of the distances between the center of the area near the P-stalk side (PS) and each aEF2 molecules, related to **c**, **d**, and **e**, respectively. In **f** and **g**, the mean ($\mu$) and SD ($\sigma$) fitted by Gaussians are 12.9 nm and 4.9 nm (**f**), respectively, and 11.5 nm and 5.2 nm (**g**), respectively. **i,** The ratios of the number of molecules of aEF2 (PS/OP), related to **c–e**.

**Supplementary Fig. 6. HS-AFM images of the hybrid 50S subunits in the presence of aEF1A and GTP.** Histogram of heights of aEF1A molecules. **b,** High-contrast sequential HS-AFM images of the hybrid 50S subunit after the addition of aEF1A•GTP on the AP-mica surface, related to Fig. 5a. aEF1A molecules bound to the ribosomal stalk are indicated by white allows. Scan area, 80 x 80 nm with 90 x 90 pixels; scan speed, 250 ms/frame; scale bar, 20 nm. **c, e,** Dot plot (left) and heat map (right) of aEF1A•GTP molecules around the hybrid 50S subunit (**c**) and aEF1A•GTP molecules around the *E. coli* 50S core (**e**), related to Fig. 5. The area near the P-stalk side (PS; radius, 20 nm) or the area opposite the P-stalk (Control; radius, 20 nm) are represented by red and blue circles, respectively. The ratios of the number of molecules of aEF1A (PS/OP) are 4.2 (**c**) and 0.9 (**d**), respectively. **d, f,** Histograms of the distances between the center of the area near the P-stalk side (PS) and each aEF1A molecules, related to **c** and **e**, respectively. In **d**, the mean ($\mu$) and SD ($\sigma$) fitted by Gaussians are 9.5 nm and 4.8 nm, respectively.

**Supplementary Fig. 7. Nucleotide dependent preferential distribution of aEF1A around the 50S subunit. a,** Typical HS-AFM images of the hybrid 50S ribosomes in the presence of 50 nM aEF1A•GDP. Scan area, 80 x 80 nm with 90 x 90 pixels; scan speed, 250 ms/frame; scale bar, 20 nm. aEF1A molecules bound to the ribosomal stalk are indicated by white arrows. **b,** Dot plot (left) and heat map (right) of aEF1A•GDP (1,915 aEF1A molecules and 211 subunits were analyzed) around the hybrid 50S subunit, related to **a**. The area near the P-stalk side (NS; radius, 20 nm) or the area opposite the P-stalk (Control; radius, 20 nm) are represented by red and blue circles, respectively. The ratios of the number of molecules of aEF1A (PS/OP) are 2.6. **c**, Histogram of the distances between the center of the area near the P-stalk side (PS) and each aEF1A molecules, related to **b**. The mean ($\mu$) and SD ($\sigma$) fitted by Gaussians are 9.5 nm and 6.4 nm, respectively. **d,** Typical HS-AFM images of the hybrid 50S ribosomes in the presence of 50 nM nucleotide free aEF1A. Scan area, 80 x 80 nm with 90 x 90 pixels; scan speed, 250 ms/frame; scale bar, 20 nm. **e,** Dot plot (left) and heat map (right) of nucleotide free aEF1A (1,088 aEF1A molecules and 89 subunits were analyzed) around the hybrid 50S subunit, related to **d**. The area near the P-stalk side (NS; radius, 20 nm) or the area opposite the P-stalk (Control; radius, 20 nm) are represented by red and blue circles, respectively. The ratio of the number of molecules of aEF1A (PS/OP) is 1.3. **f**, Histogram of the distances between the center of the area near the P-stalk side (PS) and each aEF1A molecules, related to **d**. **f,** The ratios of the number of molecules of aEF1A (PS/OP), related to Supplementary Fig. 6c, e, and Supplementary Fig. 7cb, e.

**Description of Additional Supplementary Files**

**Supplementary Movie 1**

HS-AFM movie of the *E. coli* 50S ribosomal subunits on AP-mica surface. Scan area, 60 x 50 nm with 50 x 40 pixels; scan speed, 330 ms/frame. The movie is played at 3x speed.

**Supplementary Movie 2**

HS-AFM movie of the *P. furiosus* 50S ribosomal subunits on AP-mica surface. Scan area, 160 x 160 nm with 100 x 100 pixels; scan speed, 150 ms/frame. The movie is played at 3x speed.

**Supplementary Movie 3 and 4**

HS-AFM movie of the hybrid 50S ribosomal subunits after the addition of aEF2•GTP on AP-mica surface. Scan area, 80 x 80 nm with 90 x 90 pixels; scan speed, 250 ms/frame. The movie is played at 3x speed.

**Supplementary Movie 5 and 6**

HS-AFM movie of the hybrid 50S ribosomal subunits after the addition of aEF1A•GTP on AP-mica surface. Scan area, 80 x 80 nm with 90 x 90 pixels; scan speed, 250 ms/frame. The movie is played at 3x speed.

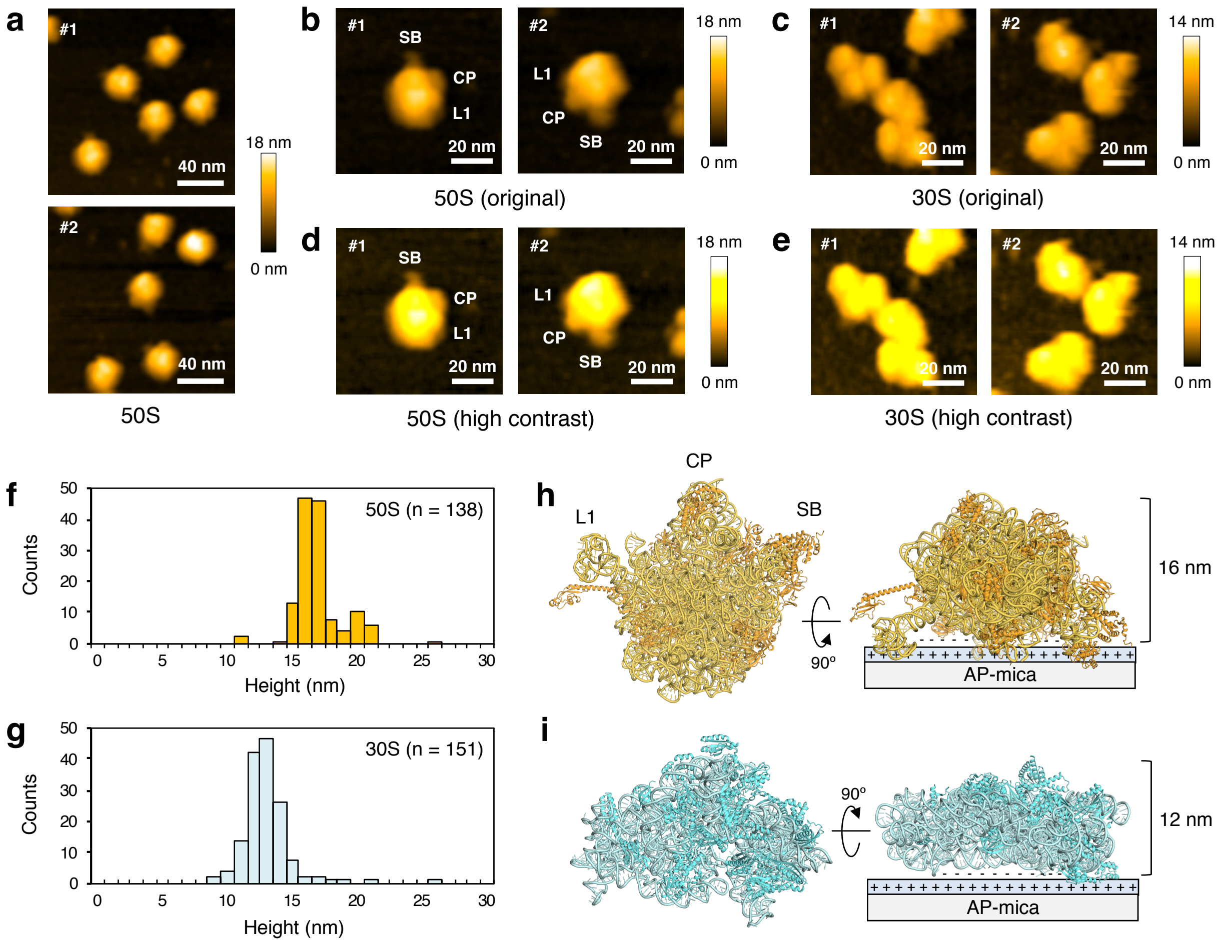


**Supplementary Fig. 1**

**a**

```
Eco     MALNLQDKQAIVAEVSEVAKGALSAVVADSRGVTVDKMTELRKAGREAG---VYMRVVRN
Tma     -MLTRQQKELIVKEMSEIFKKTSLILFADFLGFTVADLTELRSRLREKYGDGARFRVVKN
Pho     MAHVAEWKKKEVEELAKLIKSYPVIALVDVSSMPAYPLSQMRRLIRENG---GLLRVSRN
             : *:  * *:::: *      ..* ....  :::::*   **       :** :*

Eco     TLLRRAVEGTPFECLKDAFVGPTLIAYSMEH-PGAAAR--LFKEFAKAN--AKFEVKAAA
Tma     TLLNLALKNAEYEGYEEFLKGPTAVLYVTEGDPVEAVK--IIYNFYKDKKADLSRLKGGF
Pho     TLIELAIKKAAKELGKPELEKLVEYIDRGAGILVTTMNPFKLYKFLQQNRQPAPAKPGAK
        **:. *:: .  *  :  :                : .   : :* : :        ..

Eco     FEGELIPASQIDRLATLPTY---------------------------------EEAIARL
Tma     LEGKKFTAEEVENIAKLPSK---------------------------------EELYAML
Pho     VPKDVVVPAGPTPLAPGPIVGQMQALGIPARIEKGKVTIQKDTTVLKAGEVITPELANIL
        .  . . .     :*  *                                    *    *

Eco     MATMKEASAG----------KLVRTLAAVRDAKEAA------------------------
Tma     VGRVKAPITGLVFALSGILRNLVYVLNAIKEKKSE-------------------------
Pho     NALGIQPLEVGLDVLAVYEDGIVYTPDVLAIDEQEYIDMLQKAYMHAFNLAVNIAYPTPE
         .    .              :* .  .:   :.

Eco     -----------------------------------------------
Tma     -----------------------------------------------
Pho     TIEAIIQKAFLNAKTVAIEAGYITKETIQDIIGRAFRAMLLLAQQLP
```

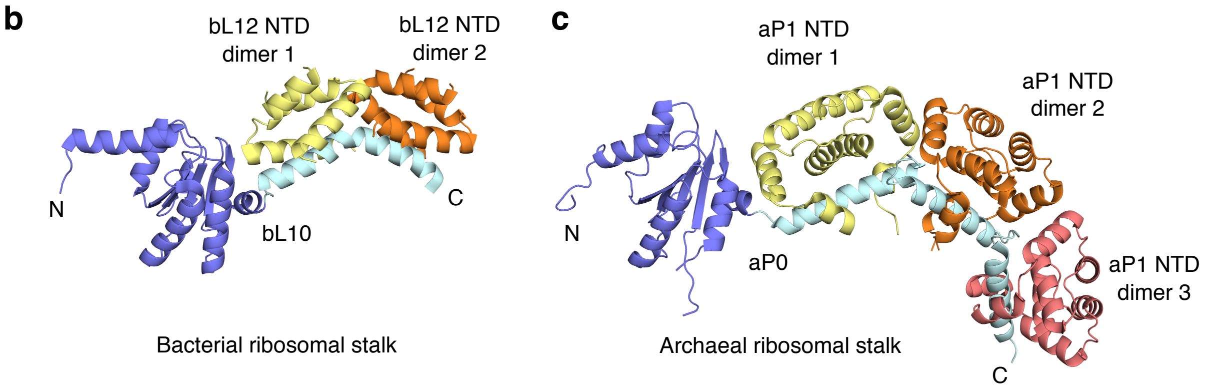


**Supplementary Fig. 2**

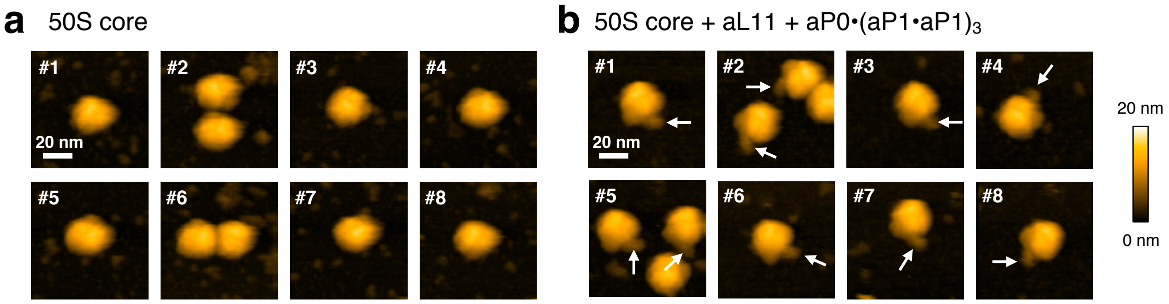


**Supplementary Fig. 3**

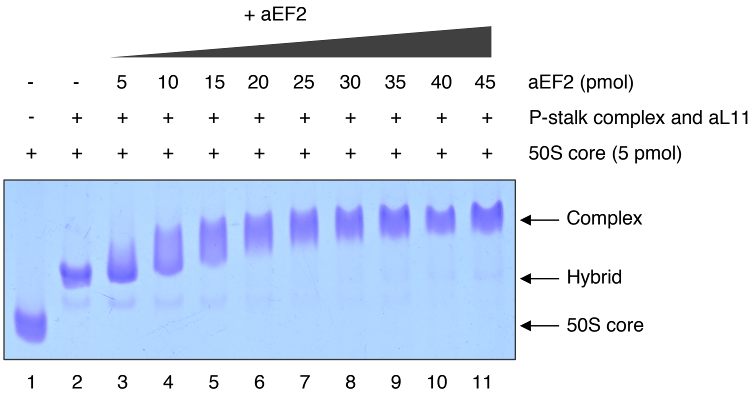


**Supplementary Fig. 4**

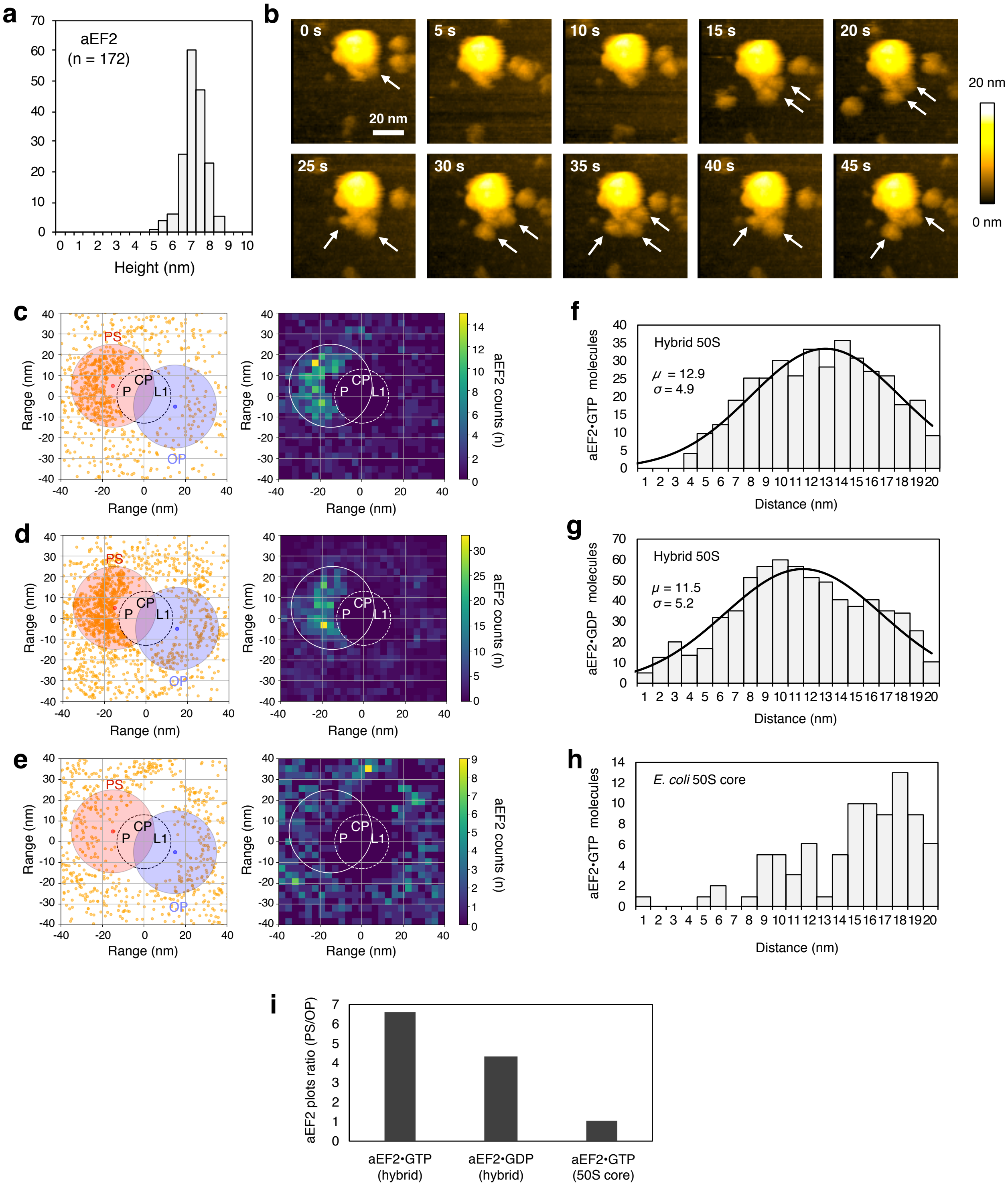


**Supplementary Fig. 5**

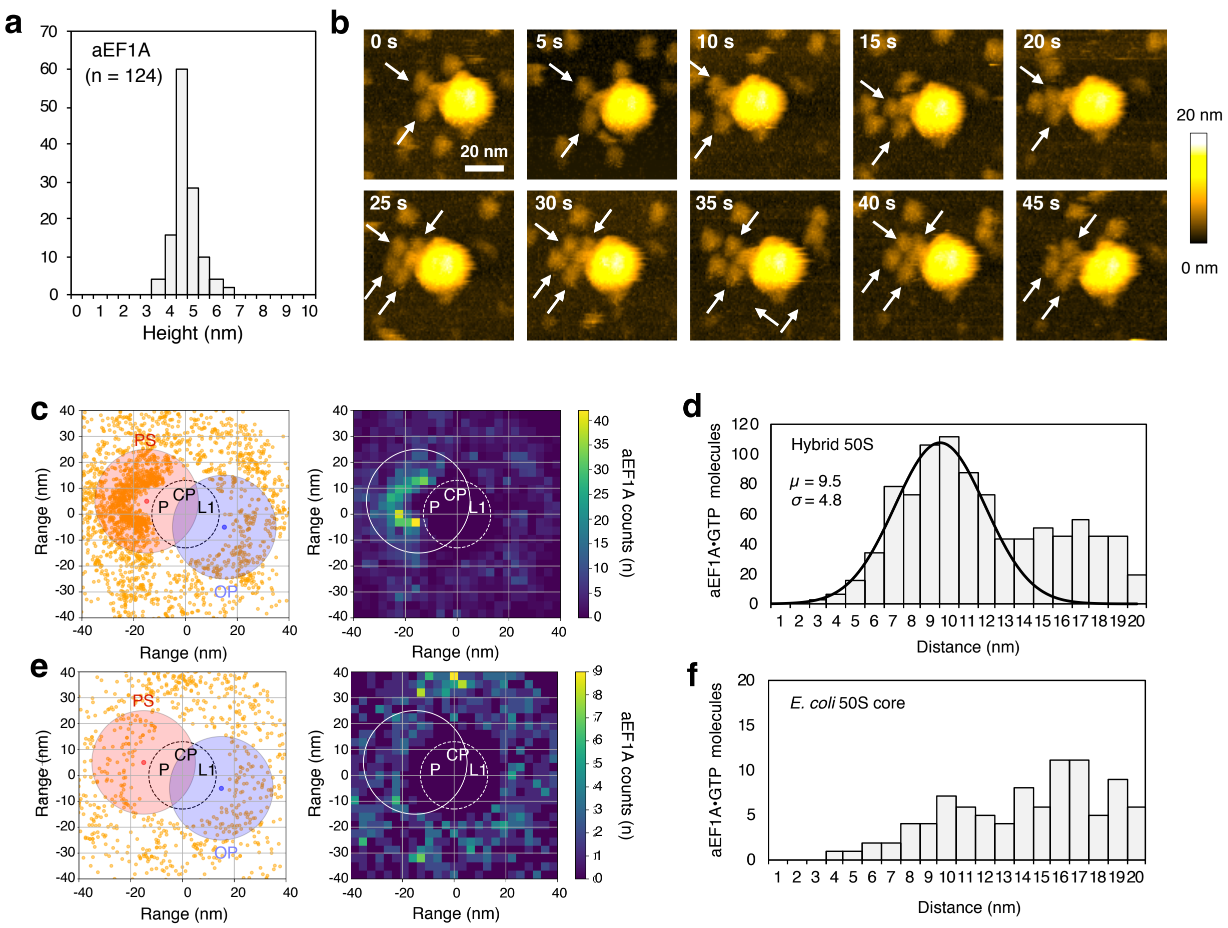


**Supplementary Fig. 6**

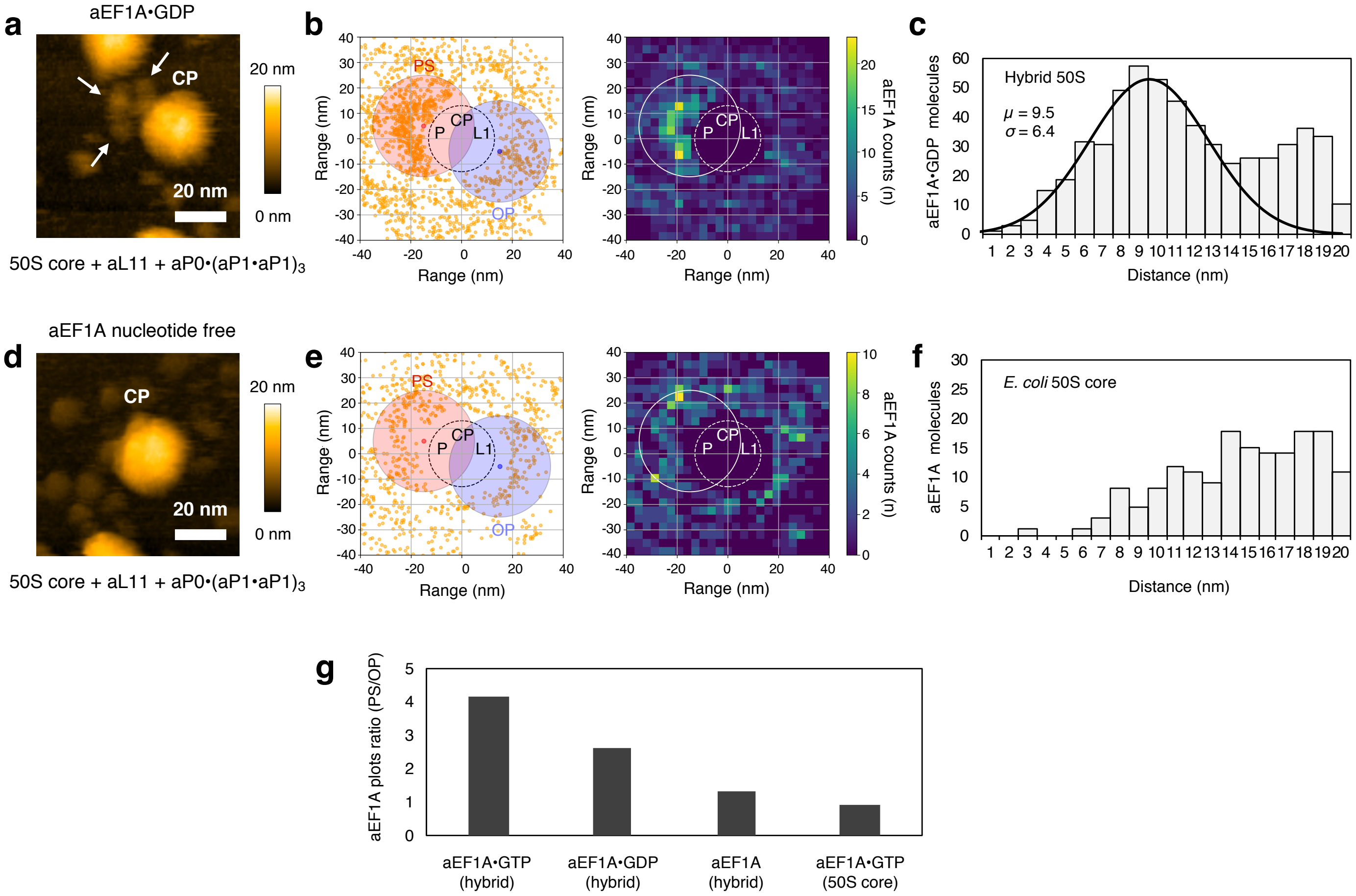


**Supplementary Fig. 7**

**Supplementary Table 1. Dissociation constant ($K_d$): aP1 C-terminus vs archaeal trGTPases**

| trGTPase | $K_d$ ($\mu$M) | Method | Reference |
|---|---|---|---|
| aEF1A•GTP | 23.6 | Fluorescent polarization | Maruyama et al. 2019 |
| aEF1A•GDP | 10.4 | Fluorescent polarization | Maruyama et al. 2019 |
| aEF2•GTP | 5.06 | SPR | Tanzawa et al. 2018 |
| aEF2•GDP | 4.05 | SPR | Tanzawa et al. 2018 |
| aABCE1 | 3.9 | Fluorescent polarization | Imai et al. 2018 |

### Supplementary Table 1
